# Which line to follow? The utility of different line-fitting methods to capture the mechanism of morphological scaling

**DOI:** 10.1101/216960

**Authors:** Alexander W. Shingleton

## Abstract

Bivariate morphological scaling relationships describe how the size of two traits co-varies among adults in a population. In as much as body shape is reflected by the relative size of various traits within the body, morphological scaling relationships capture how body shape varies with size, and therefore have been used widely as descriptors of morphological variation within and among species. Despite their extensive use, there is continuing discussion over which line-fitting method should be used to describe linear morphological scaling relationships. Here I argue that the ‘best’ line-fitting method is the one that most accurately captures the proximate developmental mechanisms that generate scaling relationships. Using mathematical modeling, I show that the ‘best’ line-fitting method depends on the pattern of variation among individuals in the developmental mechanisms that regulate trait size, and the morphological variation this pattern of developmental variation produces. For *Drosophila* traits, this pattern of variation indicates that major axis regression is the best line-fitting method. For morphological traits in other animals, however, other line-fitting methods may be more accurate. I provide a simple web-based application for researchers to explore how different line-fitting methods perform on their own morphological data.

## Introduction

The scaling relationships between body and trait sizes among adults in a species – also called static allometries – broadly captures the relative size of structures in the body and therefore characterizes species shape. Correspondingly, scaling relationships have been central to the study of morphology for well over 100 years. Early studies focused on the developmental mechanisms that underlie morphological scaling (Huxley 1924; Huxley 1932), but for the last sixty years research has largely concentrated on variation in scaling within and among species and the selective pressures that generate this variation. Morphological scaling relationships among adults in a population are typically linear on a log-log scale, and so can be modeled using the allometric equation log *y* = log *b* + *α* log *x*, where *x* and *y* are trait sizes measured in the same dimension, log *b* is the intercept and *α* is the allometric coefficient (Huxley and Tessier 1936). Log *b* broadly captures of the size of *y* relative to *x*, while *α* captures how relative trait size changes with overall size. When *α* is 1, a condition called isometry, traits scale proportionally with each other, and the size of *y* relative to *x* is maintained across body sizes. In contrast, when *α* is greater than or less than 1, called hyperallometry and hypoallometry respectively, trait *y* become disproportionally larger (*α*>1) or smaller (*α*<1) relative to trait *x* with an increase in body size. In as much as log *b* and *α* capture key aspects of body proportion, considerable effort has been invested in describing variation in log *b* and *α* among pairs of traits within a species and among species for a pair of traits.

Central to the effort to study variation in morphological scaling is the ability to fit linear relationships to morphological measurements and extract scaling parameters. This effort has been complicated by the existence of various line-fitting methods, which, when applied to the same data, can generate different slopes and intercepts. There has, unsurprisingly, been much debate over which method is ‘correct’, although no consensus has been reached (Madansky 1959; Kuhry and Marcus 1977; McArdle 1988; Warton et al. 2006; Smith 2009; Taskinen and Warton 2011; Carroll and Ruppert 2012; Hansen and Bartoszek 2012; Pelabon et al. 2013; Kilmer and Rodriguez 2016). Much of this debate has concentrated on the statistical nuances of line fitting, and in particular the assumptions different methods make about the error with which morphological measurements are made (Kilmer and Rodriguez 2016). What is often absent from the discussion is consideration of the biological phenomenon the scaling parameter is trying to capture, and the efficiency with which each method achieves this. Here, I use a well-supported developmental model of how trait size is regulated to explore which line fitting method best captures the developmental process that control the slope of morphological scaling relationships. My goal is not to declare one method of line fitting superior to another. Instead, my purpose is to encourage researchers to consider more deeply the biological processes that they are attempting to model, rather than focusing solely on the statistical aspects of the model.

## The developmental basis for morphological scaling

Morphological scaling relationships – also known as allometries – arise because there is variation in body size among individuals in a population and covariation in the size of their morphological traits (Shingleton et al. 2007). This size variation can be generated through variation in environmental factors (producing *environmental* scaling relationships) or variation in genetic factors (producing *genetic* scaling relationships). Regardless of the mechanism that generates size variation, covariation among pairs of traits arises because these traits are exposed to systemic variation in the same environmental or genetic size-regulatory factors. These factors may act directly on growing traits, for example temperature, or indirectly, for example via environmentally- or genetically-regulated growth hormones. Intuitively, it is the extent to which a change in a systemic factor generates a change in the size of each trait that determines their size covariation and consequently the slope of their scaling relationship. For example, for two traits *x* and *y*, if both traits share the same sensitivity to a size-regulatory factor, they will scale isometrically to one another as size varies with that factor (Figure 1A, B and D). In contrast, if *x* is very sensitive to changes in a size-regulatory factor but *y* is not, then as size varies in response to that factor, *x* will vary more than *y* and the slope of their scaling relationship (*x* versus *y*) will be flat (Figure 1B, C and E). Conversely, if *y* is more sensitive than *x*, then the slope of their scaling relationship will be steep. From this perspective, the slope of a morphological scaling relationship reflects the relative sensitivity of two traits to common size-regulatory factors (Shingleton et al. 2007).

**Figure 1:**
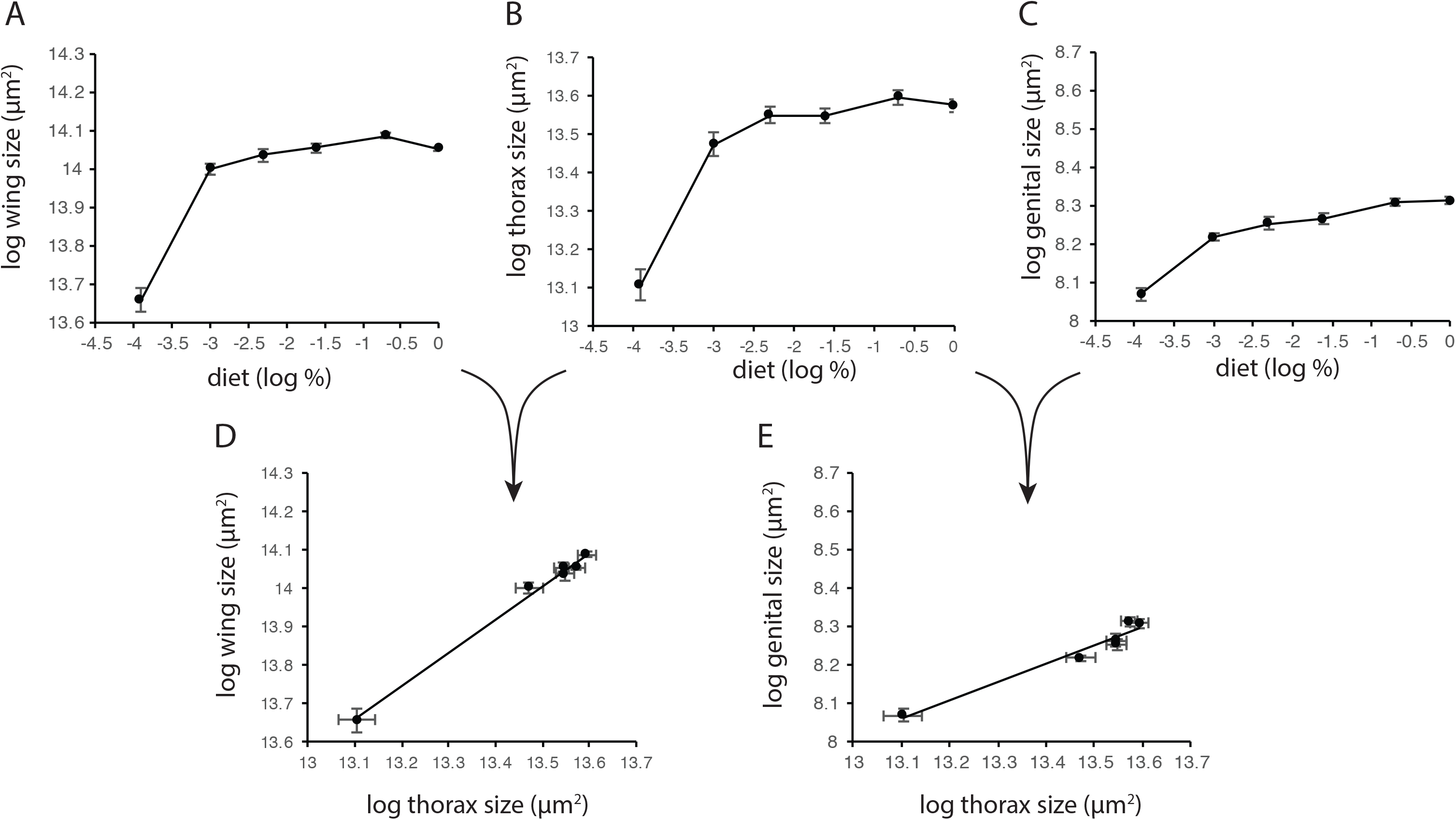
The slope of morphological scaling relationships reflects relative trait plasticity. Traits vary in the relative sensitivity to changes in developmental nutrition. In *Drosophila*, some traits, for example the wing (A) and thorax (B), are more sensitive to changes in developmental nutrition than other, for example the male genitalia (C). Consequently, the slope of the morphological scaling relationship where both traits are more nutritionally sensitive (D) is steeper than the slope where one trait is less nutritionally sensitive (E). Data from (Shingleton et al. 2009).

Empirically estimated scaling relationships are a property of a population. Trait and body size are measured for a group of individuals and fit with a line that describes how trait size changes, on average, with body size among these individuals. Consequently, scaling relationships describe the average relative sensitivity of the two traits to size-regulating factors among all individuals in the group. What is lost with this approach, however, is the relative sensitivity of the two traits for each genetically-distinct individual in a population. It is therefore useful to distinguish between an *individual-level scaling relationship*, which is the theoretical scaling relationship between adult traits for a single individual genotype across a range of potential body sizes, and the *population-level scaling relationship*, which is the observed scaling relationship between traits among individual genotypes across a range of realized body sizes (Figure 2) (Dreyer et al. 2016; O'Brien et al. 2017).

**Figure 2:**
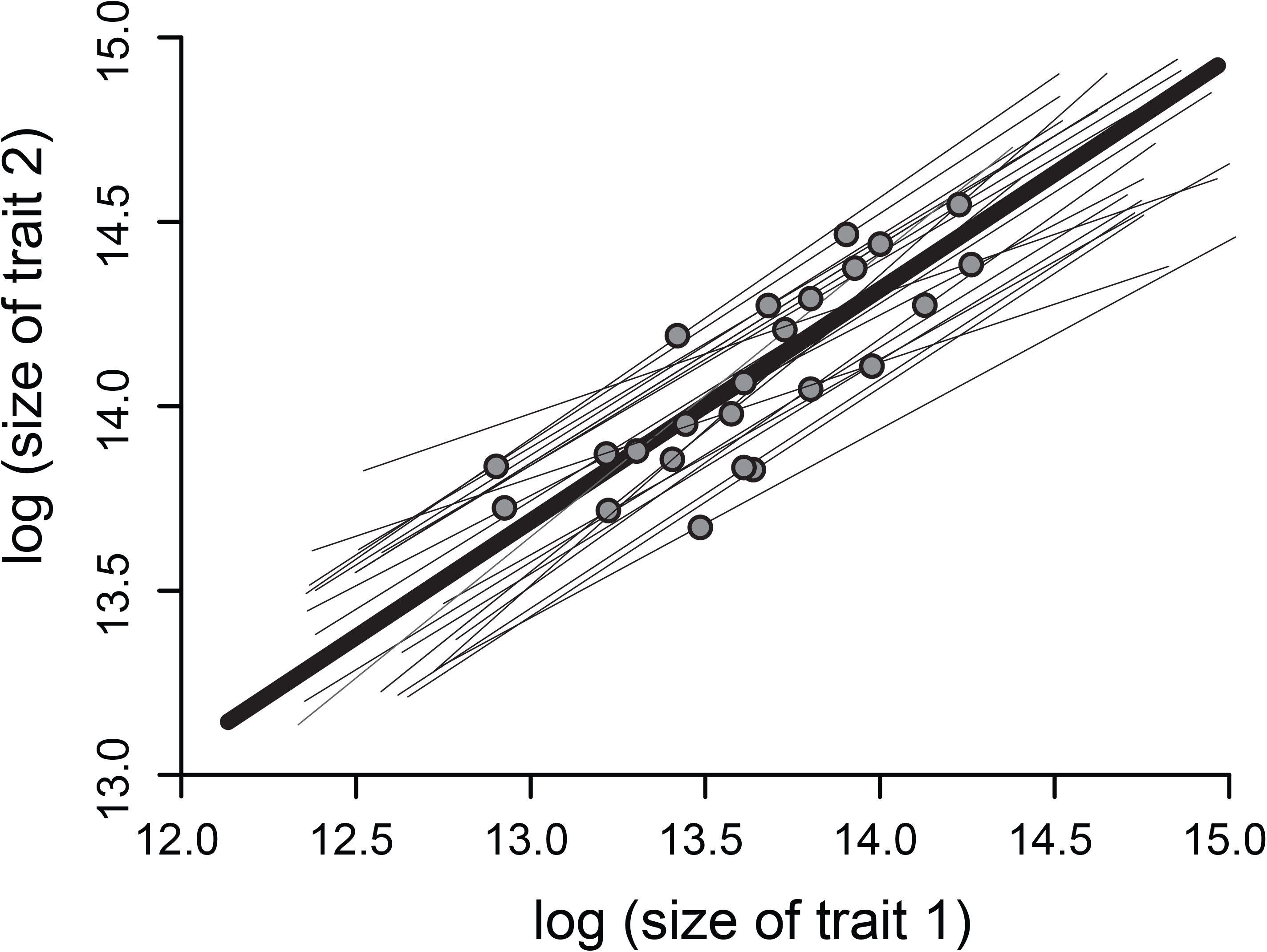
The relationship between individual and population scaling relationships. Each individual genotype will express a scaling relationship across a range of environmental conditions (for example a nutritional gradient) (thin black lines). Because each individual’s genotype is (typically) only exposed to a single developmental environment, it will sit at a single point on its cryptic individual scaling relationship. The observed population scaling relationship (thick black line) is the relationship between these individual points in the population.

Problematically, individual scaling relationships are typically unobservable, since it is not possible for the same individual to express more than one final, adult size – the one exception is where multiple individuals with the same genotype are each exposed to different levels of the same size-regulatory factor. In a genetically heterogeneous population, therefore, each individual will occupy a single point on in its otherwise cryptic individual scaling relationship (Figure 2). The observed population-level scaling relationship is that fitted among the observed phenotypes for many individuals, each a point on a cryptic individual scaling relationship. It is the variation among individuals in the relative sensitivity of traits to regulatory factors, manifest as intra-population variation in the individual-level scaling relationships, that is the ultimate target of selection on population-level scaling relationships. The pattern of this variation is thought to affect profoundly the response to selection. In as much as researchers are interested in the mechanisms that generate covariation among traits and how these mechanisms evolve, the observed population-level scaling relationship is therefore only an indirect measurement of the actual relationships of interest: the underlying cryptic individual scaling relationships.

If every member of a population had the same individual scaling relationship, and body and trait size varied only in response to systemic factors, then the population scaling relationship would be identical to all individual scaling relationships. However, in the real world, there is always scatter around a population-level scaling relationship, which has the potential to weaken the link between it and the underlying individual-level scaling relationship(s). Different line-fitting methods use different approaches to deal with this scatter. Thus, it is important to understand the cause of scatter if we are to make informed about decision about which line-fitting method to use.

There are three factors that generate scatter around the population-level scaling relationship (Figure 3). First, variation among individuals in the relative sensitivity of their traits to a systemic size-regulatory factor will generate variation in the slope of their individual scaling relationships and hence generate scatter around the observed population scaling relationship (Figure 3A & B). Second, variation in trait-autonomous size regulators will generate variation in the intercept of their individual scaling relationships, uncorrelated with variation in other traits. Such factors include genetic variation in genes expressed only in individual traits, and stochastic developmental perturbations of the sort that generate fluctuating asymmetry (Figure 3C). Third, measurement error in both traits, which, by definition, is uncorrelated among traits, will also add to scatter around the population scaling relationship (Figure 3D).

**Figure 3:**
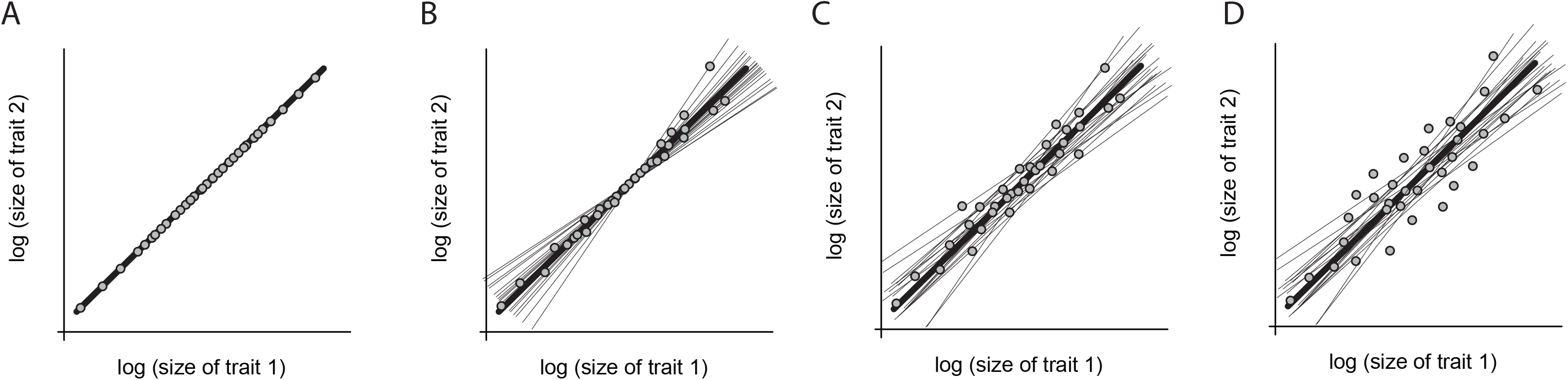
The processes that generate scatter in population scaling relationships. (A) When all individuals in a population have the same individual scaling relationship, each individual (circles) lies along the observed population scaling relationship (thick black line), which is identical to the individual scaling relationship (thin black lines, hidden). (B) When there is variation among individuals in the relative sensitivity of their trait to environmental variation, this generates variation in the slope of the individual scaling relationships (thin black lines), and generates individual scatter around the population scaling relationship. (C) When there is organ autonomous variation in trait size, this will add variation to the intercept of the individual scaling relationships, further increasing individual scatter around the population scaling relationship. (D) When there is error in measuring trait size, this will add additional individual scatter around the population scaling relationship.

## Line Fitting Methods

All regression analyses assume that there is an underlying bivariate relationship between two measurements (*x* and *y*), but that error introduces scatter around this relationship. The regression therefore attempts to fit a line through the data that minimizes the residuals; that is, the difference between an observed value and that predicted by the regression line. There are three primary methods used to fit lines to bivariate data: ordinary least squares (OLS), major axis regression (MA) and standardized major axis regression (SMA, also referred to as reduced major axis regression, RMA (Warton et al. 2006; Smith 2009; Hansen and Bartoszek 2012). These three regression methods differ in what they use as residuals, and hence what they are minimizing (McArdle 1988; Warton et al. 2006). OLS regression fits a line to bivariate data such that the vertical distance between the regression line and each point squared and summed across all points, is minimized. MA regression fits a line such that the perpendicular distance between the regression line and each point, squared and summed across all points, is minimized. The SMA regression fits a line such that the product of the vertical and horizontal distance from the line to each point, summed across all points, is minimized. The SMA is the same as the MA, but the regression line is fitted to data that are standardized so that both variables have the same standard deviation, and then rescaled back to the original axes.

The OLS is ostensibly a method for predicting the value of *y* for a known value of *x* (Montgomery et al. 2015). The slope of the OLS is the covariance between *x* and *y* divided by the variance of *x*:

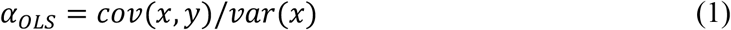

where, *var*_*x*_ is the variance of *x* and *cov*_*xy*_ is the covariance between *x* and *y*. That is, the slope of the OLS is the proportion of variation in *x* that co-varies with *y*. Implicit in OLS regression is that *x* is measured without error. Because measurement error will not affect the covariation between *x* and *y*, it follows that as measurement error for *x* increases, the denominator of Eqn 1 increases and the slope of the OLS declines. For a population-level scaling relationship, when *x* and *y* are trait sizes, this has the effect of biasing the slope down due to factors unrelated to the underlying individual scaling relationships. Further, this bias means that the slope of the OLS regression of *y* against *x* is not the inverse of the slope of the OLS regression of *x* against *y*.

The MA is the axis of the first principle component (the principle axis) calculated using the covariance matrix (Sokal and Rohlf 1995). The slope of the MA is calculated as:

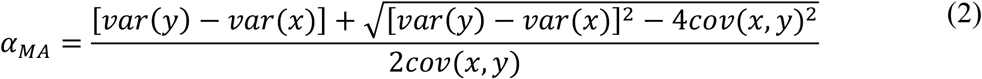

Because the MA minimizes the perpendicular distance from any point and the regression line, it assumes that deviations from the line in the *x* and *y* direction are equal: that is that the residual variance in *x* is equal to the residual variance in *y*. Implicit in this assumption is that the two traits are measured with the same error. In contrast, the SMA is the axis of first principle component calculated using the correlation matrix, and then rescaled (Smith 2009). The slope of the SMA is simply the ratio of the standard deviations of the two variables, with the sign determined by the sign of the correlation:

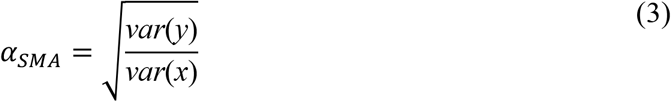

The *α*_*SMA*_ can also be calculated from *α*_*OLS*_ simply by dividing the latter by the correlation coefficient.

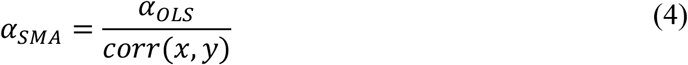

Unlike the MA, the SMA does not assume that the measurement error for the two traits are the same, but rather they have the same ratio as the variances of the two traits. Both the MA and the SMA are symmetrical, such that the slope of *x* on *y* is the inverse of the slope of *y* on *x*.

## Which Line Fitting Method to Use?

Much of the discussion about which line-fitting method should be used to estimate scaling relationships centers on the nature of the scatter around the regression line, and in particular, measurement error. Specifically, several authors have rejected the OLS method for fitting scaling relationships because it assumes that the *x* trait is measured without error, and therefore biases the slope downwards (Ricker 1973; McArdle 1988; Ebert and Russell 1994; Green 1999; Bonduriansky 2007). Others have countered this argument by demonstrating that measurement error in *x* has a marginal effect on OLS slope estimations (Hansen and Bartoszek 2012; Pelabon et al. 2013; Kilmer and Rodriguez 2016). A number of authors have observed, however, that measurement error is not the only, and unlikely to be the most important, factor that causes scatter around a regression line (Warton et al. 2006; Smith 2009; Hansen and Bartoszek 2012). As discussed above, in the case of morphological scaling relationships, scatter is also generated by variation among individuals in the relative sensitivity of traits to systemic size-regulators, and variation in trait-autonomous size-regulators. While previous authors have acknowledged the existence of this scatter, here referred to as *biological deviance*, its impact on the performance of different line-fitting methods for static allometry has not been fully explored.

Here I build a model of individual and population scaling relationships based on the developmental mechanisms that regulate trait size. I then used this model to explore how assumptions about the nature of biological deviance and measurement error impact the ability of different line-fitting methods to estimate the relative sensitivities of traits to systemic regulators of size. Other mechanisms can generate population scaling relationships – for example, genetic linkage between alleles that independently regulate the size of different traits. However, we are explicitly only interested in developmental processes that regulate trait size systemically and in a coordinated manner across the body. These are likely the primary mechanism generating population scaling relationships and will be the target of selection that generates variation in scaling among populations and species.

## A Model of Individual Morphological Scaling

The model has been published previously in a paper exploring the selection pressures that drive evolutionary changes in morphological scaling relationships (Dreyer et al. 2016). Briefly, the model assumes that trait growth is exponential and that growth rate is regulated by two factors: systemic factors, such as circulating growth hormones or temperature, and organ-autonomous factors, such as morphogens gradients within the trait. Consequently, within an individual, trait size can be modelled as:

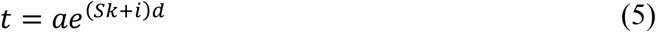

where *t* is trait size, *a* is the initial size of the trait, *S* is the level of a systemic growth factor, *k* is the sensitivity of the trait to the systemic growth factor, *i* is the organ autonomous growth rate, and *d* is the duration of growth. Log-transforming the equation generates the linear equation:

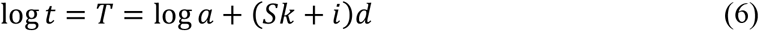

For two traits in the same body:

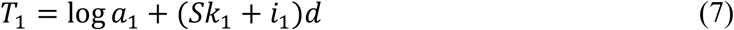

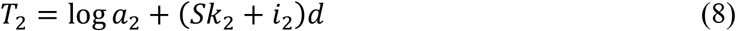

Note that when *S* is an environmental factor, Eqns 7 and 8 describe the reaction norm of trait size against the environmental variable (e.g., Figure 1A and 1B). As body size varies in response to changes in systemic growth factors, the individual scaling relationship between *T*_*1*_ and *T*_*2*_ can be described as:

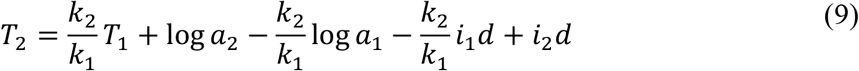

Note that the slope of the individual scaling relationship is captured by the relative sensitivity of the two traits to the systemic growth factor, *k*_2_/*k*_1_. This is supported by experimental work in *Drosophila* that demonstrates that changes in the sensitivity of a trait to changes in nutrition alters the slope of its morphological scaling relationship with other traits among genetically identical individuals, when body size varies due to diet (Tang et al. 2011; Shingleton and Tang 2012).

In a genetically heterogeneous population, individuals will vary in *k*_*1*_, *k*_*2*_, *i*_*1*_, *i*_*2*,_ *a*_*1*_, *a*_*2*_ and *d*. For simplicity, I will assume that initial trait size does not vary among individuals in a population. I will also, initially, assume that there is no variation in developmental time, although I will return to this point later in this paper. The size of individual traits therefore becomes:

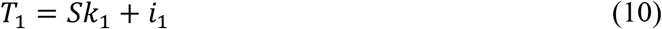

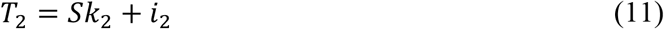

And the individual scaling relationship between *T*_*1*_ and *T*_*2*_ can be described as:

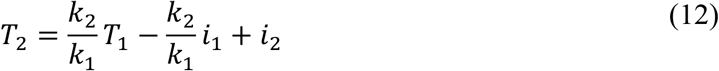

## A Model of Population Morphological Scaling

While Eqns 8 and 11 describe the scaling relationship between two traits across a range of *S* within an individual, each individual in a population is only exposed to a single level of *S* and thus only occupies a single point on this cryptic individual scaling relationship. The population-level scaling relationship between *T*_*1*_ and *T*_*2*_ estimated from multiple individuals in a population, therefore depends on population-level variation in *k*_*1*_, *k*_*2*_, *i*_*1*_, *i*_*2*_, and *S*. We can model this using Eqns 9 and 10, and by assuming that each individual’s value of *S*, *k*_*1*_, *k*_*2*_, *i*_*1*_, and *i*_*2*_ is sampled from normal distributions such that:

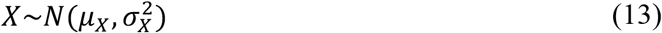

where *X* is the parameter value.

Thus, *μ*_*s*_ and 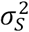 are the mean level of the systemic growth regulator, and its variation among individuals in a population, respectively; 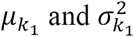 are the mean sensitivity of *trait 1* to the systemic growth regulator, and variation in this sensitivity among individuals in a population, respectively, and; 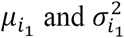 are the mean organ-autonomous growth rate of *trait 1* and variation in the organ-autonomous growth among individuals in a population, respectively. It is important to note that 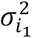 incorporates any factor that generates non-correlated variation in the size of *trait 1*, and thus includes measurement error. It is also important to note that 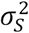 captures variation among in individuals in the level of systemic growth regulators, which may be genetic or environmental in origin. When 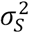 is solely a consequence of environmental variation (i.e. when all individuals are genetically identical), the population-level scaling relationship is an environmental static allometry and estimates of the individuals scaling relationship for that genotype (Shingleton et al. 2007). When 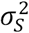 is solely a consequence of genetic variation, the population-level scaling relationship is a genetic static allometry(Shingleton et al. 2007).

The population mean of *T*_*1*_ is:

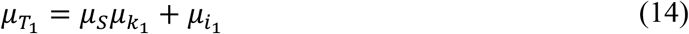

The population variance of *T*_*1*_ is:

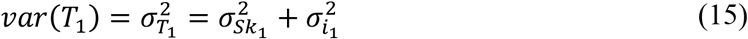

The population variance of *Sk*_*1*_ is:

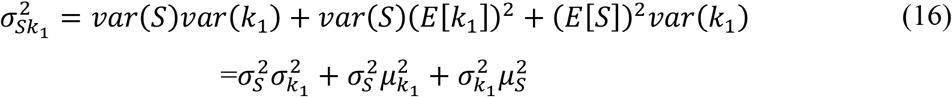

Inserting Eqn 16 into Eqn 15:

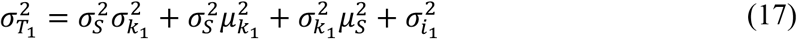

The population covariance of *T*_*1*_ and *T*_*2*_ is:

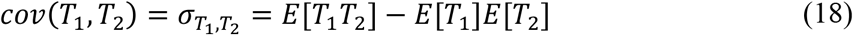

Now:

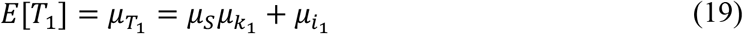

And:

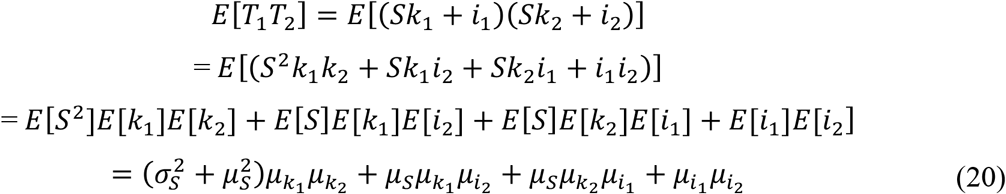

Inserting equations (17) and (18) into equation (16):

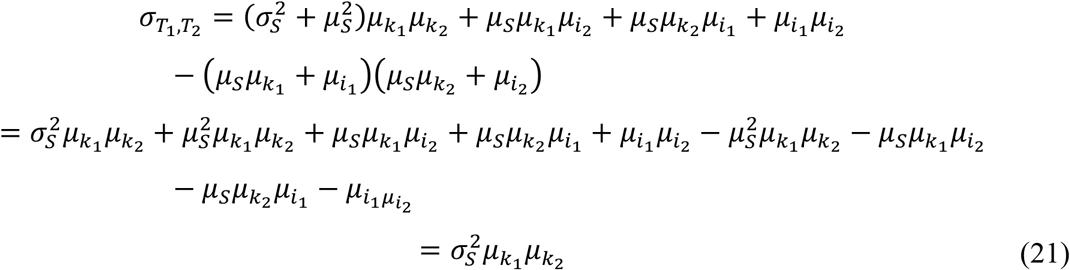

Finally, the correlation between *T*_*1*_ and *T*_*2*_ is (Sokal and Rohlf 1995):

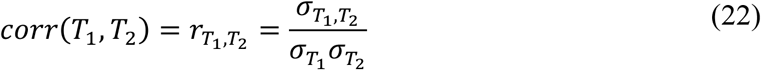

We can use the variance, covariance, and correlation of *T*_*1*_ and *T*_*2*_ to calculate the slope and intercept of the OLS, MA and SMA regressions of *T*_*1*_ against *T*_*2*_ among individuals in a population; that is, the population-level scaling relationship.

The slope of the OLS is, from Eqn 1:

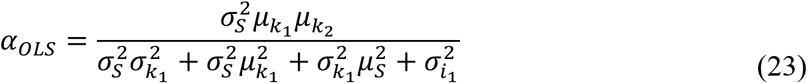

The slope of the MA is, from Eqn 2:

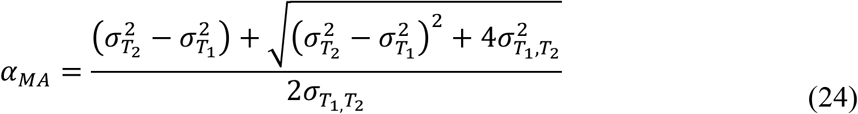

The slope of the SMA is, from Eqn 3:

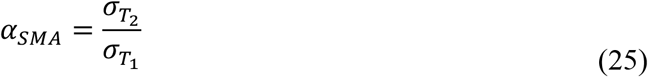

For all slopes the intercept is:

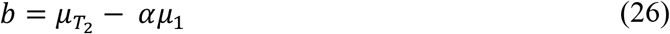

## Using the Model to Assess Line Fitting Methods

As discussed above, we are interested in the developmental processes that regulate trait size systemically and in a coordinated manner across the body and how these processes vary within and between populations and species. Consequently, we are interested in how well, and under what conditions, the different methods of calculating the population scaling relationship capture 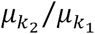-the mean relative sensitivity of traits to systemic factors that cause co-variation in trait size and that generate scaling relationships.

For the OLS method, re-arranging Eqn 23 with respect to 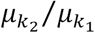:

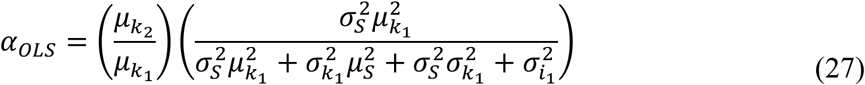

Thus, the slope of the OLS is 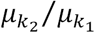 multiplied by a bias factor that is between 0 and 1. It follows that 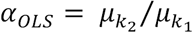 when 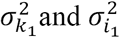 are zero. As these parameters increase *α*_*OLS*_ becomes smaller relative to 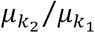. Conversely, as 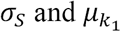 increase, *α*_*OLS*_ approaches 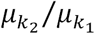.

For the MA slope, re-arranging Eqn 24 with respect to 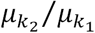:

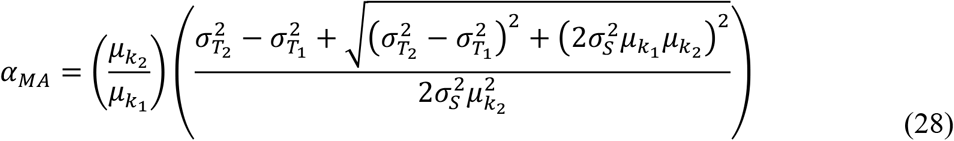

Again, the slope of the MA is 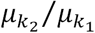 multiplied by a bias factor. Here, the bias is unbound and can be greater or less than 1. However, when 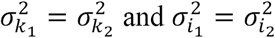, the bias factor reduces to one and 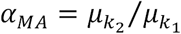 (Supplementary Material). Thus, unlike the OLS, the MA slope can capture perfectly the average relative sensitivity of traits to systemic factors that cause covariation in trait size, albeit under certain conditions.

For the SMA method, re-arranging Eqn 25 with respect to 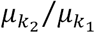:

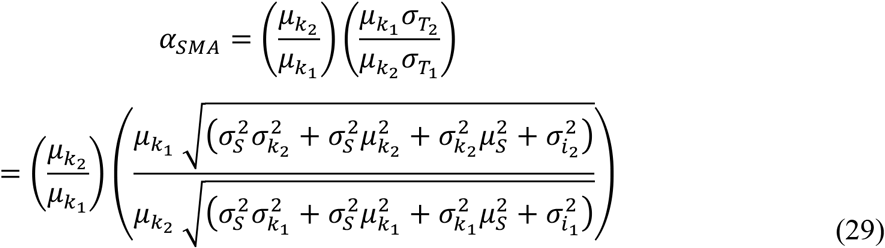

As for the MA slope, the bias of the SMA slope can be greater or less than 1, but will equal 1 when 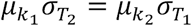: that is, if the ratio of the trait standard deviations 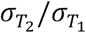 equals 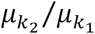. The SMA slope will also be unbiased when 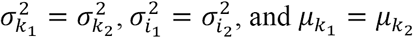. Thus, like the MA slope, and unlike the OLS slope, the SMA slope can also capture perfectly the average relative sensitivity of traits to systemic factors that cause co-variation in trait size.

While both the SMA and MA can capture 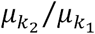, the conditions under which they do so may be biologically restrictive; both 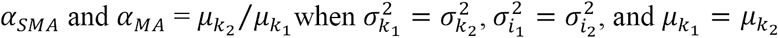. That is, both methods will capture the mean individual-level scaling relationship between two traits when the traits scale (on average) isometrically, and when they show the same level of organ-autonomous size variation and variation in sensitivity to systemic growth regulators. Below I examine how reasonable these conditions are.

For many animals, body proportion appears to be largely maintained across a range of body sizes, suggesting that most traits scale near-isometrically and 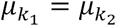. However, this may not be the case; a recent meta-analysis of 553 static allometries indicated that for traits not obviously subject to sexual selection, the average slope is 0.87 (95% CI: .7*9-.94)* (Voje 2016). This suggests that slight hypoallometry is the most common scaling relationship. One caveat is that many, if not all, of these slopes were calculated using an OLS against body size. This will lower the estimate of the slope whenever there is uncorrelated variation in whatever proxy of body size was used, either due to measurement error or because of variation in organ-autonomous size-regulators, or whenever there is variation in the sensitivity of body size to systemic size-regulators. Thus, it is not clear if slight to moderate hypoallometry is the norm. Further, even if most traits scaled isometrically, much of the research on allometry concentrates on traits that most obviously deviate from isometry, for example the hyperallometric secondary sexual characteristics of stalk-eyed flies (Wilkinson 1993) or horned beetle (Warren et al. 2013), or the hypoallometric genitalia of male arthropods (Eberhard 2009). For these traits 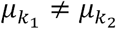.

Both the SMA and the MA can, however, capture 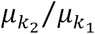 even when 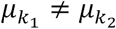. For the SMA, it can be seen from Eqn. 29 that as 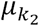 and the slope increase, the standard deviation of 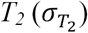 will also increase but at a slower rate, biasing the slope of the SMA down. Under these conditions, the SMA slope will only capture 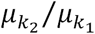 if the other factors that contribute to 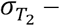 that is, 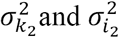 – increase also. From a biological perspective, this would imply that the mechanisms that regulate *k,* the sensitivity of a trait to systemic growth regulators, also regulate the population-level variance of *k* and of *i*, the organ-autonomous growth rate. While it is conceivable that the same mechanism could regulate the mean and variance of *k* (e.g.(Emlen et al. 2012)), it is difficult to envision how this mechanism could also regulate the variance of *i*, which by definition acts organ-autonomously. In contrast, the MA can capture 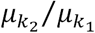 when 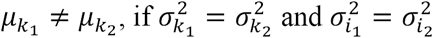. This makes intuitive sense. The MA assumes that residual variance in *x* is equal to the residual variance in *y*, which will be true if traits share the same variation in sensitivity to systemic growth regulators, and have the same level of organ-autonomous variation in growth rate. While we know the developmental mechanisms that regulate a trait’s sensitivity to at least one systemic growth regulator (insulin-like peptides) (Tang et al. 2011), and have elucidated many of the developmental mechanisms that regulate organ autonomous growth (Irvine and Harvey 2015), there has been no study to directly measure genetic variation in these mechanisms regarding trait size. However, as I discuss below, it is possible to get an idea of what this level of variation is, at least indirectly.

## Estimating Parameter Values from Data

While, to my knowledge, no one has explicitly measured the level of genetic variation in the sensitivity of organs to changes in systemic growth regulators, nor genetic variation in organ-autonomous growth, there are published data that can be used to estimate these values. Specifically, Dreyer and Shingleton (Dreyer and Shingleton 2011) measured the size of the wing, the femur of the first leg, maxillary palp, posterior lobe of the genital arch and anal plate of 50 males from 40 isogenic *Drosophila* lineages. The static allometry among trait sizes within a lineage is the cryptic individual static allometry (Eqn 12), while the static allometry of mean trait sizes among lineages is the population static allometry (Eqn 23-26).

It is possible to use these data to estimate genetic variation in sensitivity to environmentally-regulated growth regulators, that is, a gene-environment (GxE) interaction. Within an isogenic lineage, all size variation is due to environmental variation and measurement error. For each lineage, the slope of a bivariate allometry is the ratio of *k*_*2*_ to *k*_*1*_, and variation in *k*_*2*_ to *k*_*1*_ leads to variation in the slope. Problematically, the ratio of two normally-distributed variables does not have a well-defined variance (unlike the product of two normally-distributed variables), and so it is not straightforward to calculate the 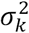 for the two traits from the variance of their ratios. However, for each lineage it is possible to calculate the relative plasticity of each trait against the plasticity of overall body size, using multivariate allometric coefficients. These are the loadings of the first eigenvector of the variance-covariance matrix for trait size, when trait size varies in response to environmental factors, which it does in an isogenic line of flies. Multiplying the loadings by 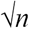, where *n* is the number of traits measured, gives the plasticity of a trait relative to overall body size. The relative plasticity of a trait is a measure of its sensitivity to environmentally-regulated systemic growth regulators. The variation in relative trait plasticity among lineages can therefore be used as a measure of variation in this sensitivity. Figure 4 shows this variation, which is not significantly different among traits (Brown-Forsythe Test, *P* = 0.6458 (Feltz and Miller 1996)).

**Figure 4:**
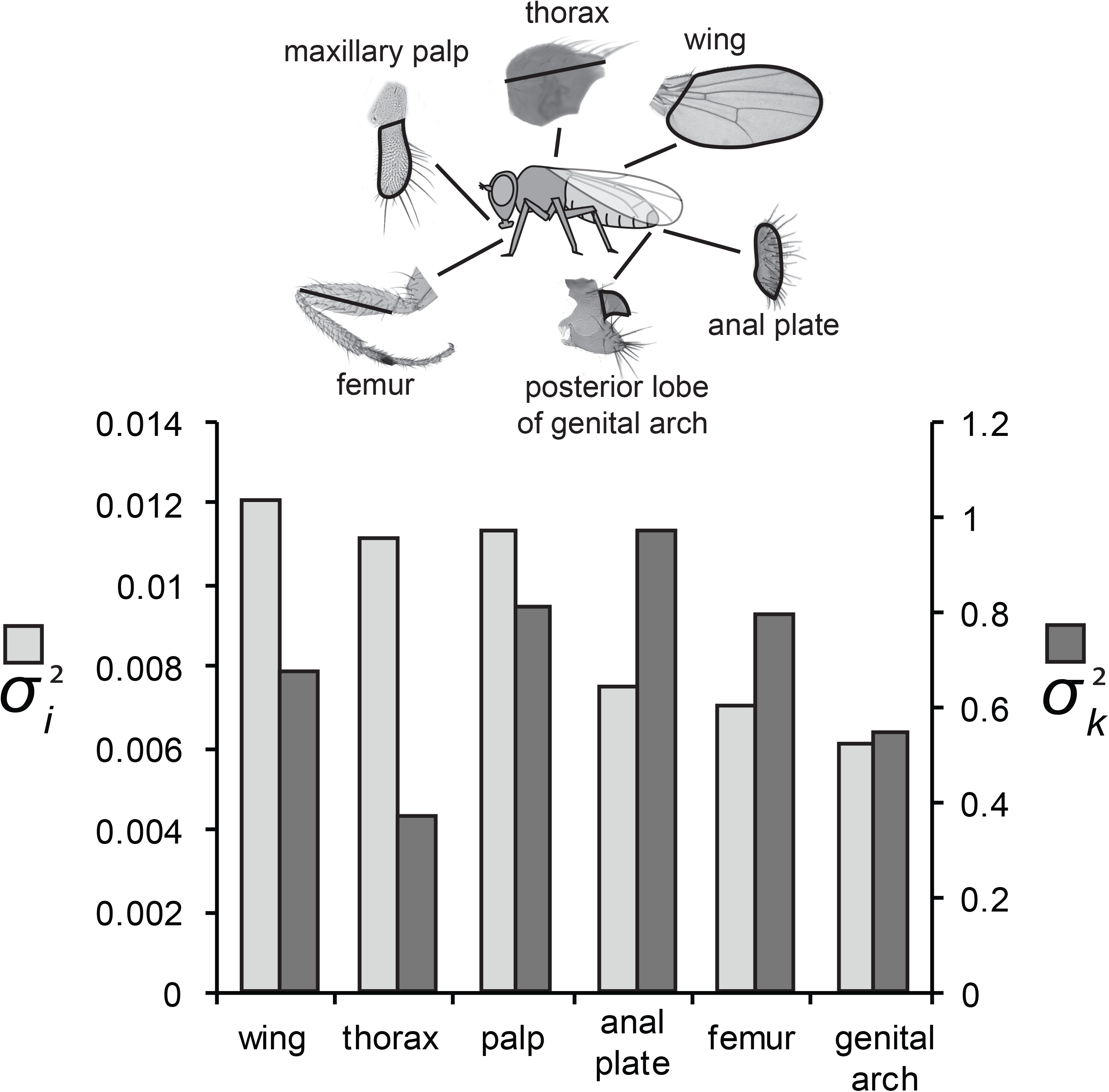
Estimates of *σ*_*i*_ (light gray bars, left axis) and *σ*_*k*_ (dark gray bars, right axis) for six morphological traits in *Drosophila*. Neither *σ*_*i*_ nor *σ*_*k*_ varied significantly among traits ((Brown-Forsythe Test, *P* > 0.6458 for both). Illustration shows morphological traits measured. Data from (Dreyer and Shingleton 2011).

It is not possible, however, to do the same for genetically-regulated systemic growth regulators, that is the gene-gene (GxG) interaction. In theory, *k* for a genetically-regulated systemic growth regulator can be calculated among individuals that are genetically identical except at the locus that controls the level of this regulator. Such ‘co-isogenic lineages’ would be challenging to generate, even in tractable model organisms such as *Drosophila*. Further, this would then have to be repeated among multiple co-isogenic lineages with different genetic backgrounds to estimate 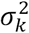. Nevertheless, it is reasonable to assume that most of the genetic variation in systemic growth factors is a consequence of genetic variation in the synthesis and release of hormones that are also environmentally co-regulated, for example insulin-like-peptides and growth hormone. Consequently, the variance of relative trait plasticity may be a good approximation of 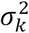 for genetically-regulated systemic growth regulators.

The same data can be used to calculate *μ*_*i*_ for each trait and hence *σ*_*i*_ among lineages. If we assume that the cryptic individual scaling relationship for a lineage is described by Eqn 12, the intercept of the scaling relationship between two traits, *C*_*1,2*_, can be calculated as 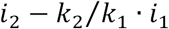, where *k*_2_/*k*_1_ is the slope of the scaling relationship. We can therefore use the individual scaling relationship of *trait 1* against *trait 2*, of *trait 2* against *trait 3,* and *trait 3* against *trait 1*, to construct a system of three linear equations:

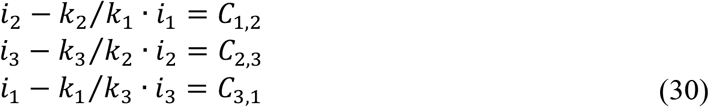

Solving these equations using the observed values for the slopes (*k*_*y*_/*k*_*x*_) and intercepts (*C*_*x,y*_) of the individual scaling relationships (fitted using a linear regression) gives us the values of *i*_*1*_, *i*_*2*_ and *i*_*3*_ for the lineage. This methodology can be expanded to calculate *i* for any number of traits. These values can then be used to calculate 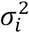 for each trait among lineages. Figure 5 shows this variation for each *Drosophila* trait, which is not significantly different among traits (Brown-Forsythe Test, *P* =0.8141).

**Figure 5:**
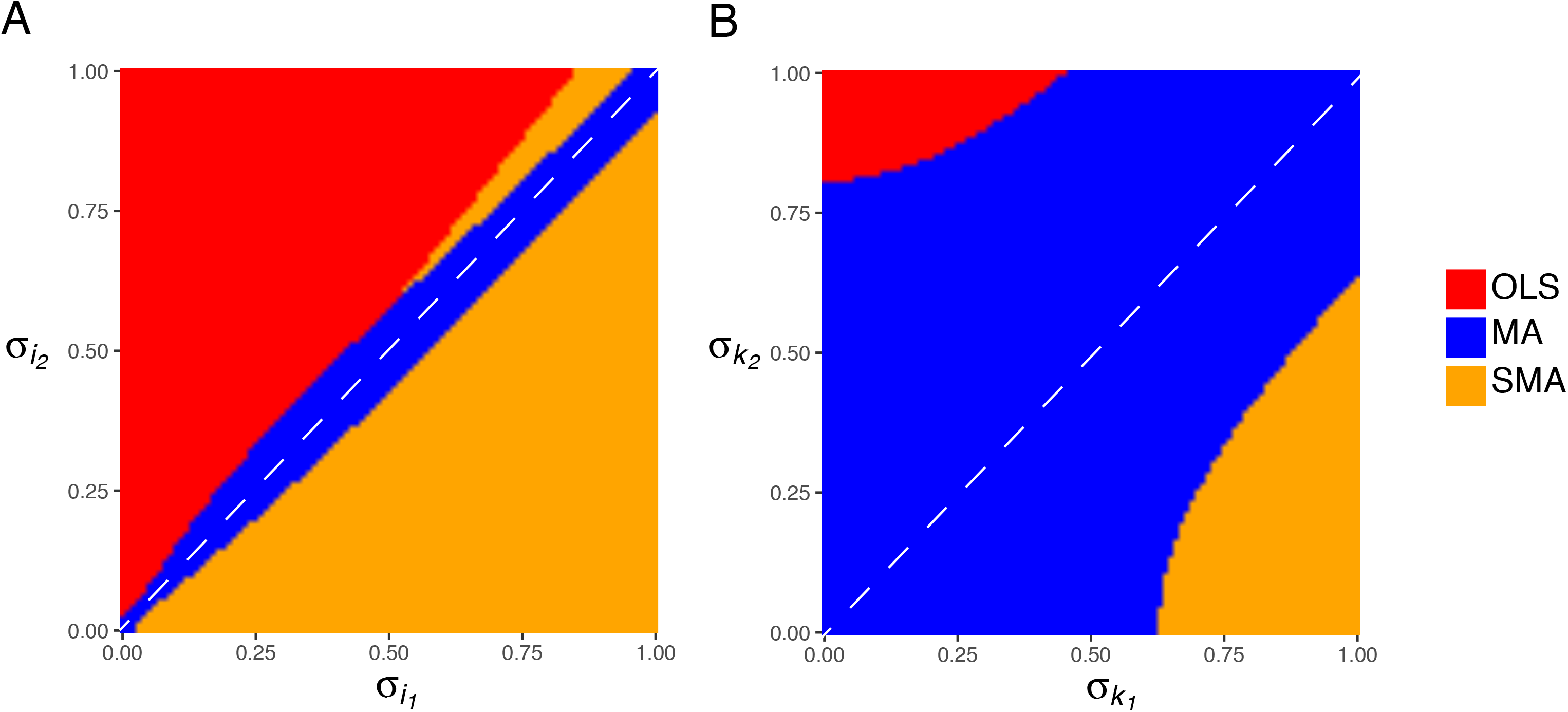
The influence of different aspects of trait variation on the effectiveness of different line-fitting methods to capture the slope of morphological scaling relationships. At each point, the method (red:OLS; blue: MA; orange: SMA) that generates a slope closest to the slope of the average individual scaling relationship 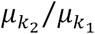 for the population is displayed. (A) The influence of variation in trait sensitivity to systemic growth regulators, *σ*_*k*_, on the effectiveness of different line-fitting methods. (B) The influence of organ-autonomous variation in growth rate, *σ*_*i*_, on the effectiveness of different line-fitting methods.

Collectively, therefore, published morphological data in *Drosophila* suggest that traits do not vary significantly in 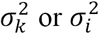, supporting the application of MA regression to estimate 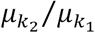 — the mean relative sensitivity of traits to systemic factors that cause co-variation in trait size and that generate scaling relationships.

## Exploring the Parameter Space

While data from *Drosophila* may support the application of MA regression to calculate the slope of allometric relationships, the same need not be true for other traits in other organisms. I therefore developed an interactive interface using *shinyapp* that allows a user to determine which line fitting method (OLS, SMA, MA) best captures the average slope of individual scaling relationships 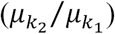 in a population under different conditions. The application can be accessed at https://shingletonlab.shinyapps.io/linefitting/. Alternatively, the *R*-scripts that run the application are available on Dryad, which a user can download and run locally on their computer.

The interface of the application is shown in Figure 3. Briefly, the application allows the user to explore the how well different line-fitting methods fit simulated data across a range of model parameter values. At the bottom of the interface, the user can assign parameter values to the model. The user can then select one of three plots to explore the effect of the parameter values on the utility of the different line-fitting methods. The interface is described in more detail in the Supplementary Material. The application assumes the ‘best’ line fitting method is the one that produces a slope closest to 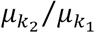. Unsurprisingly, this depends on the parameters used to generate the population scaling relationship. Nevertheless, there are two general trends that are worth highlighting:

First, the model suggests that organ-autonomous variation in growth rate (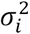) for the two traits has the greatest influence on which line fitting method best captures 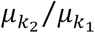 (Figure 4A). As outlined above, the MA perfectly captures 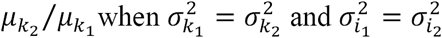. However, when 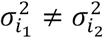, which is the best line-fitting method depends on whether the underlying relationship is hyperallometric 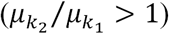 or hypoallometric 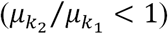. For hypoallometric scaling relationships, the SMA tends to be best line fitting methods when 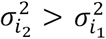, while the OLS tends to be the best line-fitting method when 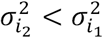. In contrast, for hyperallometric relationships, the SMA tends to be the best line-fitting method regardless of whether 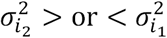. When 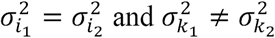, the MA still best captures 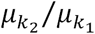 unless the population scaling-relationship is nearly isometric (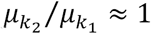), in which case the SMA appears to be the best method (Figure 4B).

Second, the means of the parameter values have comparatively little influence on how well each line fitting method captures 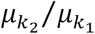. Specifically, *μ*_*i*_ has no influence on the regression slope using any line fitting method, while *μ*_*s*_ affects only the OLS and SMA regression slope, and only marginally. As discussed above, both 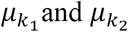 influence how well different line fitting methods capture the slope, but also regulate the slope itself.

## Conclusions and Future Directions

The goal of this study was to examine the performance of different line fitting methods in capturing the mechanisms that produce covariation in trait size and generate morphological scaling relationships within populations. The model highlights that the phenotype most morphological researchers measure when studying scaling relationships – the population-level scaling relationship - is an imperfect representation of the relationship they are, in many cases, implicitly most interested in - the individual-level scaling relationship. The observed population-level scaling relationship is not generated by a ‘true’ scaling relationship, with individual scatter around this relationship being a consequence of observation error or stochastic biological processes. Rather, the population-level scaling relationship is the observable part of a population of cryptic individual-level scaling relationships. The key insight is that departure from the population-level scaling relationship is, in part, due to variation among the slopes of individual-level scaling relationships for the group. It is this variation that evolution acts upon to generate changes in allometry. Explicitly incorporating this variation into the model of morphological scaling not only allows one to better understand how scaling evolves, but also what statistical methods one should use to detect when evolution has occurred. Finally, this study has quantified in *Drosophila* the level of genetic variation in key developmental parameters that regulate morphological scaling, finding no evidence for differences among traits in their variance.

This is certainly not the first study to explore which line-fitting method should be employed to model morphological static allometries. However, most earlier studies did not fully consider the sources of variation that generates scatter around scaling relationships. One important exception is the work of Hansen and Bartoszek (2012) who applied a similar model to explore the interplay between biological and measurement error in evolutionary regressions, including evolutionary scaling relationships (that is, morphological scaling relationships among species). As with this study, they started with the premise that all line-fitting methods impose bias. However, they concluded that the bias imposed by OLS regression is less severe than that imposed by MA and SMA regression, and therefore favored the OLS method of line-fitting to evolutionary and allometric regressions. Their model did not, however, consider the sources of biological variation that generate scatter in population scaling relationships. Nevertheless, both Hansen & Bartsoszek’s and this study reiterate the importance of considering the sources of variation when applying regression models to biological data.

Which method the reader uses will depend on the purpose of their regression and the levels of variation that generate scatter in their morphological scaling relationship. If the reader is interested in estimating the mean relative sensitivity of traits to systemic size-regulators that generate covariation in size (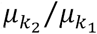), then our model suggests that the ‘best’ method for modeling morphological scaling relationships is most dependent on the level of organ-autonomous variation in trait size, and the level of error in measuring those traits, *σ*_*i*_. In contrast, the level of variation in sensitivity of a trait to systemic size-regulators, *σ*_*k*_, has relatively little effect on the ability of different line-fitting methods to capture 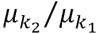. For morphological traits in *Drosophila*, the MA regression appears to be the best method, although this is based on a relatively small data set, where there was relatively little variation in trait size within isogenic lineages. Thus, the observed lack of heteroscedasticity for *i* and *k* among traits, may reflect the low power of the Brown-Forsythe test for equality of variance. Consequently, I am being deliberately agnostic on the question of which line-fitting method is best. There are few published data that allow full assessment of this question. What is needed are more data revealing how variable are cryptic individual scaling relationships within populations.

Finally, it is important to note that the insight provided by the model is limited by its veracity. The model has the advantage of being very simple, and assumes that traits have linear ‘reaction norms’ in response to variation in systemic regulators of size (Eqn 10 and 11). (Here, ‘reaction norm’ includes trait response to systemic factors that may be genetic). For many organismal traits, this will not be true. Nevertheless, even if the trait ‘reaction norms’ are not linear, they can still generate linear individual scaling relationships (Figure 1). For example, if

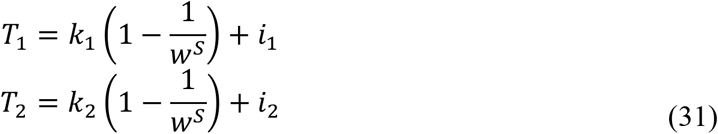

where *T*_*1*_ and *T*_*2*_ are the (log) sizes for two traits, *S* is the systemic growth regulator, *k*_*1*_ and *k*_*2*_ is the total variation in trait size from *T*_*min*_ to *T*_*max*_, *w* is a factor that controls the rate at which *T*_*1*_ and *T*_*2*_ approaches their asymptote, and *i*_*1*_ and *i*_*2*_ are the size of *T*_*1*_ and *T*_*2*_ due to intrinsic growth. As long as *w* is the same for both traits, the relationship between *T*_*1*_ and *T*_*2*_ is described by Eqn 12, and the slope of the scaling relationship is controlled by *k*_*1*_/*k*_*2*_.

Including developmental time makes the model more complex. If developmental time is variable, but the same for both traits and unaffected by the systemic growth regulator, then the individual scaling relationship is described by:

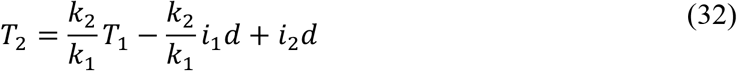

where *d* is developmental time. Here variation in developmental time acts in the same way as variation in trait-autonomous growth, with the same effect on the efficiency of the different line-methods to capture 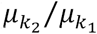. If, however, the systemic growth regulator also affects growth duration trait-autonomously, then the model becomes more complex. Such a model is detailed in the Supplementary Material and indicates that the slope of the morphological scaling relationship is controlled by the sensitivity of a trait’s growth rate and growth duration to changes in the systemic size regulator.

It is possible to use any growth model to explore how variation in underlying growth parameters affect the efficiency of different line fitting methods to capture the values of those parameters. Needless to say, the more complex the developmental model, the more difficult it is to mathematically describe the slopes and intercepts of the population scaling relationship (Eqn 23-25). Nevertheless, even with the most complex models of individual growth it is trivial to generate a simulated population scaling relationship *in silico*, and explore how changes in model parameters affect the slopes and intercept of the population-level scaling relationship when fit using different line-fitting methods. In general, there are likely multiple developmental mechanisms that generate co-variation in trait size among individuals in a population, and a corresponding number of models. As we learn more of these mechanisms, our statistical methods should be adapted to better capture their key characteristics. It is these mechanisms, after all, that are ultimately the target of selection for changes in morphological scaling.

## Acknowledgments

AWS was supported by National Science Foundation grant nos. IOS-1557638, IOS- 1406547 and IOS-0919855, and by Lake Forest College. The author thanks Tony Frankino for his comments on early versions of the manuscript and Enrique Treviño for providing mathematical advice.

---

**Supplementary Figure 1:** Screen shot of the application that allows users to explore the effect model parameters on the ability of different line-fitting methods to capture the slope of morphological scaling relationships. The first tab shows the population scaling relationship for two traits in a population of 500 individuals. The red, blue, and orange line shows the theoretical (sold line) and sampled (broken line) OLS, MA and SMA regression. The green line shows the average individual scaling relationship 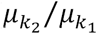 for the population. The application is available at: https://shingletonlab.shinyapps.io/linefitting/

